# The midpoint of cortical thinning between late childhood and early adulthood differs between individuals and brain regions: Evidence from longitudinal modelling in a 12-wave neuroimaging sample

**DOI:** 10.1101/2022.02.10.479868

**Authors:** Delia Fuhrmann, Kathrine Skak Madsen, Louise Baruël Johansen, William FC Baaré, Rogier A Kievit

**Affiliations:** Kings College London; Danish Research Centre for Magnetic Resonance, Copenhagen University Hospital Hvidovre; Danish Research Centre for Magnetic Resonance, Centre for Functional and Diagnostic Imaging and Research, Copenhagen University Hospital; Cognitive Neuroscience Department, Donders Institute for Brain, Cognition, and Behavior, Radboud University Medical Center

**Keywords:** brain maturation, adolescence, cortical thickness, nonlinear mixed models, sex differences, rostral middle frontal gyrus, rostral anterior cingulate

## Abstract

Charting human brain maturation between childhood and adulthood is a fundamental prerequisite for understanding the rapid biological and psychological changes during human development. Two barriers have precluded the quantification of maturational trajectories: demands on data and demands on estimation. Using high-temporal resolution neuroimaging data of up to 12-waves in the HUBU cohort (*N* = 90, aged 7-21 years) we investigate changes in apparent cortical thickness across childhood and adolescence. Fitting a four-parameter logistic nonlinear random effects mixed model, we quantified the characteristic, s-shaped, trajectory of cortical thinning in adolescence. This approach yields biologically meaningful parameters, including the midpoint of cortical thinning (MCT), which corresponds to the age at which the cortex shows most rapid thinning - in our sample occurring, on average, at 14 years of age. These results show that, given suitable data and models, cortical maturation can be quantified with precision for each individual and brain region.

## Introduction

The human cortex undergoes protracted microscopic and macroscopic structural changes between childhood and adulthood^1^. Individual differences in cortical structure have been associated with a range of phenotypic differences, including physical and mental health^2–5^, neurodevelopmental disorders as well as cognitive performance in childhood and adolescence^2,6^. Although other measures of brain structural development such as brain volume or white matter connectivity provide complementary insights into brain maturation, cortical thinning is one of the most widely used proxies of brain maturation^7–9^. We here use the term *apparent* cortical thickness throughout to highlight that Magnetic Resonance Imaging (MRI) studies measure a proxy of cortical thickness, that follows a similar spatial patten to that observed in histological studies^10^, but has absolute values that may be influenced by signal intensities and contrasts^11^.

Previous work has generally observed developmental decreases in cortical thickness from childhood (after 2-3 years of age) to early adulthood^8^, with longitudinal studies showing that the rate of cortical thinning increases in adolescence^9,12^. There is an emerging consensus on a characteristic s-shaped, non-linear trajectory at the population level (Figure 1)^9,12^. This process of cortical thinning is thought to reflect a range of underlying biological processes^13^, including increasing myelination of the deeper cortical layers^14^ and decreasing synaptic density^15^.

**Figure 1.**
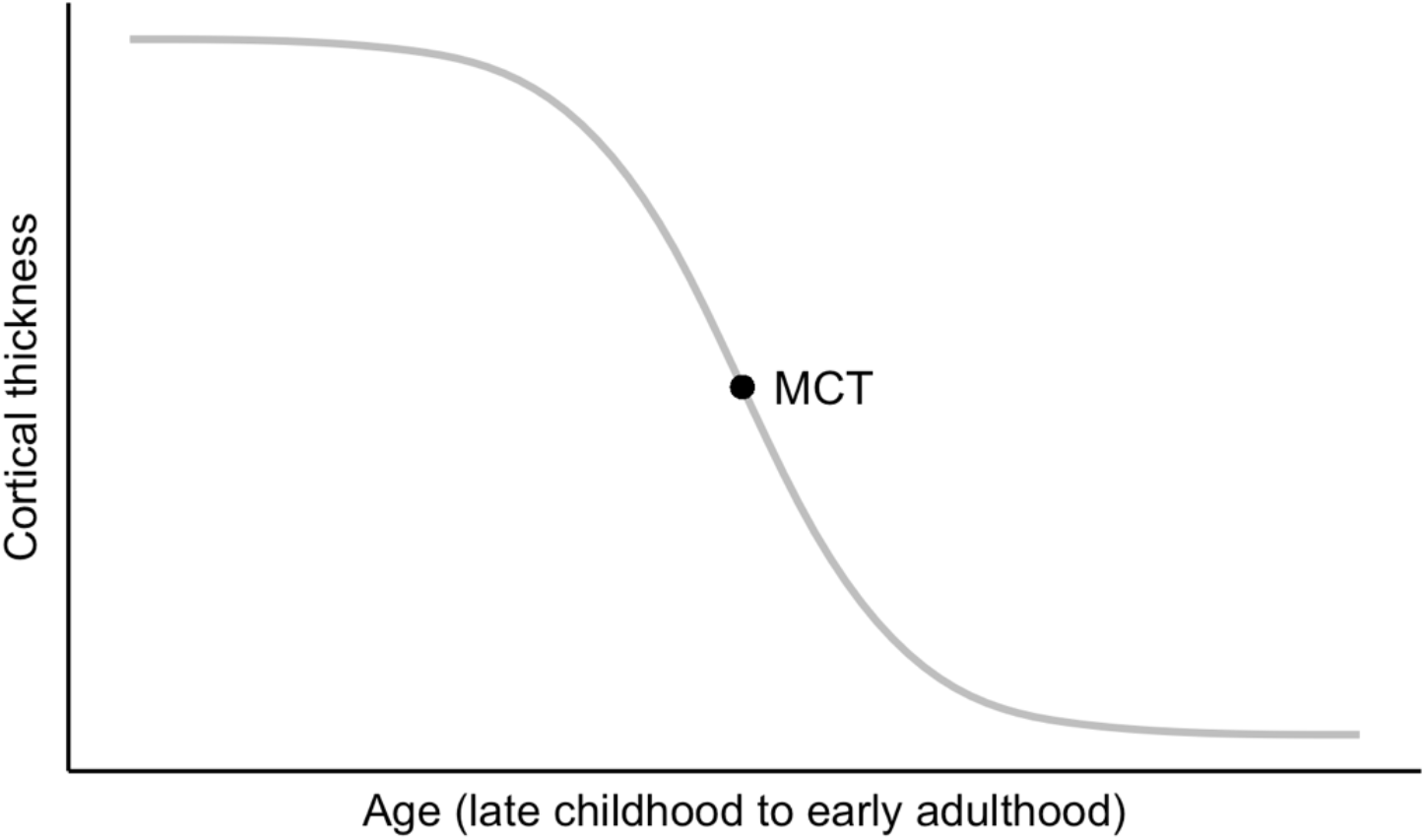
Schematic illustration of cortical thinning during adolescence, showing the Midpoint of Cortical Thinning (MCT), i.e., the age of most rapid cortical thinning.

Hypotheses concerning individual differences in the process of cortical thinning and their link to psychological development feature prominently in neurodevelopmental theories. For instance, it has been hypothesized that a life history of adversity may lead to accelerated^16,17^ or delayed^18,19^ cortical maturation. The developmental mismatch theory suggests that a mismatch in maturity between subcortical and cortical brain regions (with frontal regions commonly thought to thin last) may help explain the prevalence of risk-taking behaviour during adolescence when this maturation disparity is thought to be maximal^20,21^. At the group comparison level, hypotheses posit that girls demonstrate earlier cortical maturation than boys^22–24^, with these differences hypothesized to underlie developmental differences in behavioural and psychopathological phenotypes. Similarly, Nunes et al. (2020) hypothesized that children with autism spectrum disorder are characterized by accelerated brain maturation^25^. Overall, hypothesized differences in cortical maturation are central to some of the most influential neurodevelopmental theories. However, the data and analytic methods currently used to capture maturation are no match for ambitions in understanding and applying the construct of cortical maturation.

Longitudinal data is costly and time-consuming to procure, therefore, most neuroimaging studies to-date rely on cross-sectional data. Cross-sectional data precludes the investigation of developmental changes and cross-sectional measures necessarily conflates distinct sources of cortical thickness differences (baseline thickness, the onset of maturation, as well as speed and total amount of thinning). Only under extremely restrictive assumptions (e.g., identical brain thickness in early childhood and late adolescence, identical rates of thinning) can a cross-sectional measure be used as a proxy for development. Given that these assumptions are known to be empirically untrue^1^, our empirical knowledge of cortical maturation is likely to be extremely limited.

Where longitudinal data does exist, the average number of waves per subject is generally below three, and the time between scans is 2.5 years on average^26^. This limits our ability to observe more subtle brain changes in time periods with ongoing maturation such as adolescence, as well as our ability to capture the trajectory of cortical thinning, which is known to be non-linear^27^. This is particularly true at the individual level. That said, large cohorts with multiple time points, like ABCD^28^, are currently emerging.

Most longitudinal studies investigating individual differences to date have used linear modelling approaches, such as linear mixed effects models, using the percentage of, or absolute change, in cortical thickness between two ages as an indicator of maturation of a given brain region, with larger changes commonly equated to more protracted maturation^29,30^. However, these estimates are typically confounded by the initial thickness of a region - thicker regions can thin more than thinner regions. Even when such confounds are controlled for, the absolute change in thickness remains dependent on the precise age range studied. Alternative, nonlinear approaches, such as Generalized Additive Mixed Models (GAMMs)^31^, can capture complex nonlinear relationships and are excellent tools for predictive purposes. Nonlinear mixed models offer a similarly flexible approach for capturing complex nonlinear relationships to GAMMs. A particular advantage of nonlinear mixed models is that they yield readily interpretable parameters, informative, of, e.g., ages of rapid development (Figure 1), making them an attractive, but currently underused, tool for developmental neuroscientists.

To address the challenge of quantifying cortical maturation at the individual level, we here leverage a unique dataset (with up to 12 longitudinal measurements between late childhood and early adulthood) and a quantitative framework currently underutilized in cognitive neuroscience (non-linear random effects modelling) to demonstrate that cortical maturity can be defined, and estimated, at the individual level, offering a new window of insight into cortical development across adolescence. Nonlinear mixed models are a powerful tool that can capture developmental processes at the individual level^31^ and yield readily interpretable parameters. One of these parameters allows us to provide a novel quantitative definition of cortical maturation during adolescence at the individual level: The *midpoint of cortical thinning* (MCT, see Figure 1). The MCT is the point in adolescent development where the rate of cortical thinning is at its peak for an individual. We show that the MCT can be used to study the extent to which cortical maturation differs between individuals, sexes, and brain regions.

## Materials and Methods

### Cohort

The present longitudinal study included data from 90 typically developing children and adolescents (53 females, 37 males), who were enrolled in the longitudinal HUBU (“*Hjernens Udvikling hos Børn og Unge*”, Brain Maturation in Children and Adolescents) study. The HUBU study was initiated in 2007, where 95 participants (55 females, 40 males) aged 7 - 13 years and their families were recruited from three elementary schools in the Copenhagen, DK, suburban area^32^. All children whose families volunteered were included, except for children with a known history of neurological or psychiatric disorders or significant brain injury. Prior to participation, all children assented to the study procedures and informed written consent was obtained from parents. Informed written consent was also obtained from the participants themselves when they turned 18 years of age. The study was approved by the Ethical Committees of the Capital Region of Denmark (H-KF-01-131/03 and H-3-2013-037) and performed in accordance with the Declaration of Helsinki. Participants were scanned up to 12 times with scanning intervals of 6 months for the first 10 visits, one year between visits 10 and 11, and three years between visits 11 and 12 (Figure 2).

**Figure 2:**
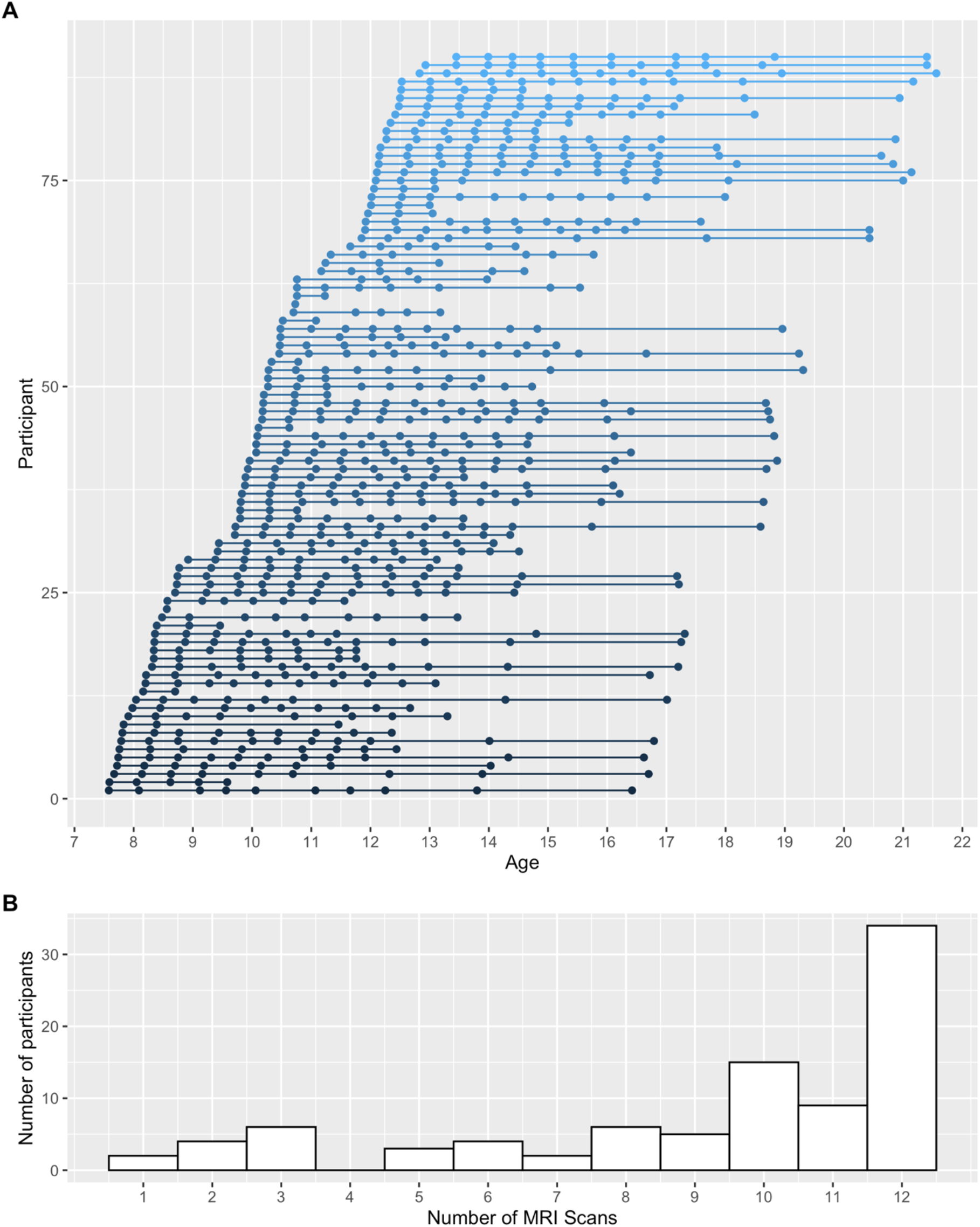
Spacing and timing of scans for each participant (Panel A) and a histogram of the number of scans included in the analysis (Panel B).

Here, we included data from the first 12 assessments of the HUBU study. All baseline MRI scans were evaluated by an experienced neuroradiologist, and all raw images were visually inspected to ensure sufficient quality. Five participants were excluded from the present study, due to receiving a psychiatric diagnosis after study initiation (*N* = 2), incidental clinical finding on the MRI scan (*N* = 1), or no MRI scans or FreeSurfer outcomes of sufficient quality (*N* = 2). Our final sample analysed here consisted of 90 participants (53 females, 37 males) aged 7.6 - 21.6 years. For this sample, we excluded 73 MRI sessions if one of the following criteria was met: participants did not finish the MRI session (2 participants, 2 scans), participants were not scanned due to metallic dental braces (14 participants, 31 scans), had poor MR-image quality (22 participants, 31 scans), or had acquired a brain injury after baseline (1 participant, 9 scans). A total of 745 valid MRI scans (scans per participant: range = 1 - 12, mean = 8.3, median = 10, interquartile range (*IQR*) = 8 – 12) were included in the statistical analyses. Data from the HUBU cohort has previously been used in cross-sectional^33–37^ and longitudinal^32,38^ studies examining brain-behavioural relationships.

### MRI protocol

Participants underwent structural MRI on a 3T Siemens Magnetom Trio MR scanner (Siemens, Erlangen, Germany) using an eight-channel head coil (Invivo, FL, USA). Two T1-weighted images were acquired using a 3D MPRAGE sequence (TR = 1550 ms, TE = 3.04 ms, matrix = 256 × 256, 192 sagittal slices, 1 × 1 × 1 mm^3^ voxels, acquisition time = 6:38). A T2-weighted image was acquired using a 3D turbo spin echo sequence (TR = 3000 ms, TE = 354 ms, FOV = 282 × 216, matrix = 256 × 196, 192 sagittal slices, 1 × 1 × 1 mm^3^ voxels, acquisition time = 8:29).

### FreeSurfer pre-processing and extraction of cortical thickness

All T1-weighted and T2-weighted images were processed using tools available in the FreeSurfer (version 6.0) software suite^**39–41**^. Cortical surface reconstruction was implemented using the following procedures: skull stripping, non-uniformity correction, white matter segmentation, creation of initial mesh, correction of topological defects, and creation of optimal white and pial surfaces^**39–41**^. Images were then processed with the longitudinal stream^**42**^ in FreeSurfer, to estimate changes in cortical thickness across time. Apparent cortical thickness was calculated as a measure of the shortest distance between the white and pial surfaces. Cortical grey matter parcellations were based on surface-based nonlinear registration to the Desikan-Killiany atlas based on gyral and sulcal patterns and Bayesian classification rules^**41**^, yielding estimates for 34 ROIs in each hemisphere. Cortical thickness estimates were averaged across the hemispheres.

To check the quality of the FreeSurfer outputs, we followed the Enigma protocol (http://enigma.ini.usc.edu/protocols/imaging-protocols/). We used statistical outlier detection for average thickness and surface area of each cortical parcel. Statistical detection of both within-and between-subject outliers was performed based on the age, sex, and age-by-sex adjusted residuals derived using GAMMs, to account for age and sex differences in the brain measures. Scans of concern were then visually inspected. In line with the Enigma protocol, we did not perform manual editing on the FreeSurfer outputs and instead discarded data of questionable quality. Based on these quality checks, we excluded one participant from the entire study. Furthermore, we excluded the following cortical parcels from statistical analysis for *N* participants: right temporal pole (*N* = 1), left frontal pole (*N* = 1), right paracentral (*N* = 1), right cuneus (*N* = 1), right lateral occipital (*N* = 1), and left fusiform gyrus (*N* = 2).

Finally, the Euler number, which indicates the topological complexity of the reconstructed cortical surface, was extracted from FreeSurfer for each scan and used as a proxy for in-scanner motion and a quantitative measure of image data quality in our statistical analyses, to account for potential systematical bias in data quality across age. The Euler number has been correlated with visual image quality inspection scores as well as with cortical thickness in a regionally heterogeneous pattern across datasets^**43**^. The Euler number was extracted for each hemisphere and summed to produce one value per scan.

### Statistical analyses

We here modelled cortical thinning between childhood and adulthood using nonlinear mixed models implemented via the saemix^44^ package (version 2.4) in R (version 4.1.0) and RStudio (version 1.4.1717). To capture the characteristic s-shape of cortical thinning, we fit the four-parameter logistic function^45^, as defined in the Results section. See [BLINDED] for our analysis code. Cortical thickness was modelled as the dependent variable and age as the independent variable. We fit a model to mean cortical thickness, as well as for one for each of the 34 ROIs of the Desikan-Killiany atlas^41^. We included sex at birth and in-scanner motion as covariates in all models, to account for potential differences thereof in developmental trajectories^46,47^. In-scanner motion was operationalized by the Euler number (*M* = -137.07, *SE* = 5.33, range = -298.2 - -69.08). Females were coded as the reference group. While sex was a covariate of interest, and motion a covariate of no interest, both are modelled the same way in the NLMM framework. Covariate parameters can, in principle, be included to control for the effect of motion and sex on any of the main fixed effect parameters of the model (in our case, the two asymptotes, the MCT, and the hill). This yields a maximum of eight covariates. To avoid potential overfitting with a highly parametrized model^48^, we determined the optimal covariate model via forward stepwise model selection using mean thickness as the outcome variable. To the best of our knowledge, no consensus methodology for model selection with covariates exists for nonlinear mixed models. We use forward selection as a pragmatic and tractable solution in the context of demanding model estimation. Although some have advocated for more complex model selection algorithms, such as best subset selection^49^, recent work suggests that forward selection can perform similarly to these alternatives^50^. The process starts with no covariate parameters in the model. It then iteratively adds the variable that best improves the fit. For all covariate parameters not in the model (e.g., eight at the first iteration), we check their *p*-value when added to the model one at a time. We chose the covariate parameter with the lowest *p*-value less than 0.05. This process was continued until no new predictors could be added. Based on this model selection process, we allowed the upper and lower asymptote, and MCTs to differ between sexes and included the motion parameter as a covariate for the lower asymptote (see Supplementary Table 1 for the full results). The remaining parameters were held constant for sex and motion. Estimates were obtained for the four parameters of the logistic function, as well as the two covariates. Precision was assessed by inspecting the coefficient of variation (CV) for each parameter, as provided by saemix. CVs are standardized measures of dispersion and are calculated as the ratio of the standard deviation to the mean. CVs < 20% are generally considered acceptable^51^.

In a second step, we assessed differences in MCT estimates across different brain regions. We assessed whether MCTs across brain regions could be constrained to equality using Confirmatory Factor Analysis (CFA), as implemented in lavaan^52^ for R (version 0.6-9). We used full information maximum likelihood with robust standard errors to account for missingness and nonnormality. We estimated a one-factor CFA model in which factor loadings were freely estimated for each brain region. We then compared this model to one where the factor loadings were still freely estimated, but the intercepts were constrained to equality using the likelihood ratio test. A significant likelihood ratio test here indicates a loss in model fit for a constrained model, providing evidence that brain regions differ in their MCTs.

Third, we used Exploratory Factor Analysis to assess the dimensionality of MCTs across regions and to identify maturational factors capturing developmental trends across regions. This was implemented through parallel analysis via the psych^53^ package for R (version 2.1.6), using an oblique oblimin transformation.

Finally, we tested the popular hypothesis of a posterior-anterior gradient of development^54^ by correlating MCTs for each region with their y-coordinates in MNI space. Coordinates were obtained from the brainGraph^55^ package in R (version 3.0.0). In an additional exploratory analysis we also correlated MCTs for each region with their z-coordinates in MNI space to test for a potential dorsal-ventral gradient.

## Results

Cortical thickness showed nonlinear changes between childhood and early adulthood across cortical regions (Figure 3). The characteristic sigmoid, or s-shaped, curve, of cortical thinning, was apparent across most brain regions. We captured this shape using the four-parameter logistic function:

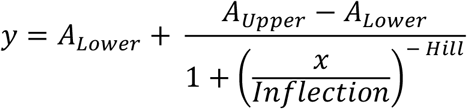

The function yields four biologically meaningful parameters: The upper asymptote (*A*_*Upper*_, maximal apparent thickness in mm), lower asymptote (*A*_*Lower*_, minimal apparent thickness in mm), and *Hill*, the slope of change, and the *Inflection*. We were particularly interested in the latter parameter, which corresponds to the MCT, and was used here as an index of cortical maturation (see Figure 1). The MCT can be compared across individuals, sexes, and brain regions. To this end, we first modelled cortical thickness averaged across the cortex, identified the average pattern of maturation, and assessed sex differences. Next, we modelled cortical thickness for different brain regions to investigate patterns of maturation across the cortex. Sex and in-scanner motion, as a proxy for image quality, was controlled for in all analyses.

**Figure 3.**
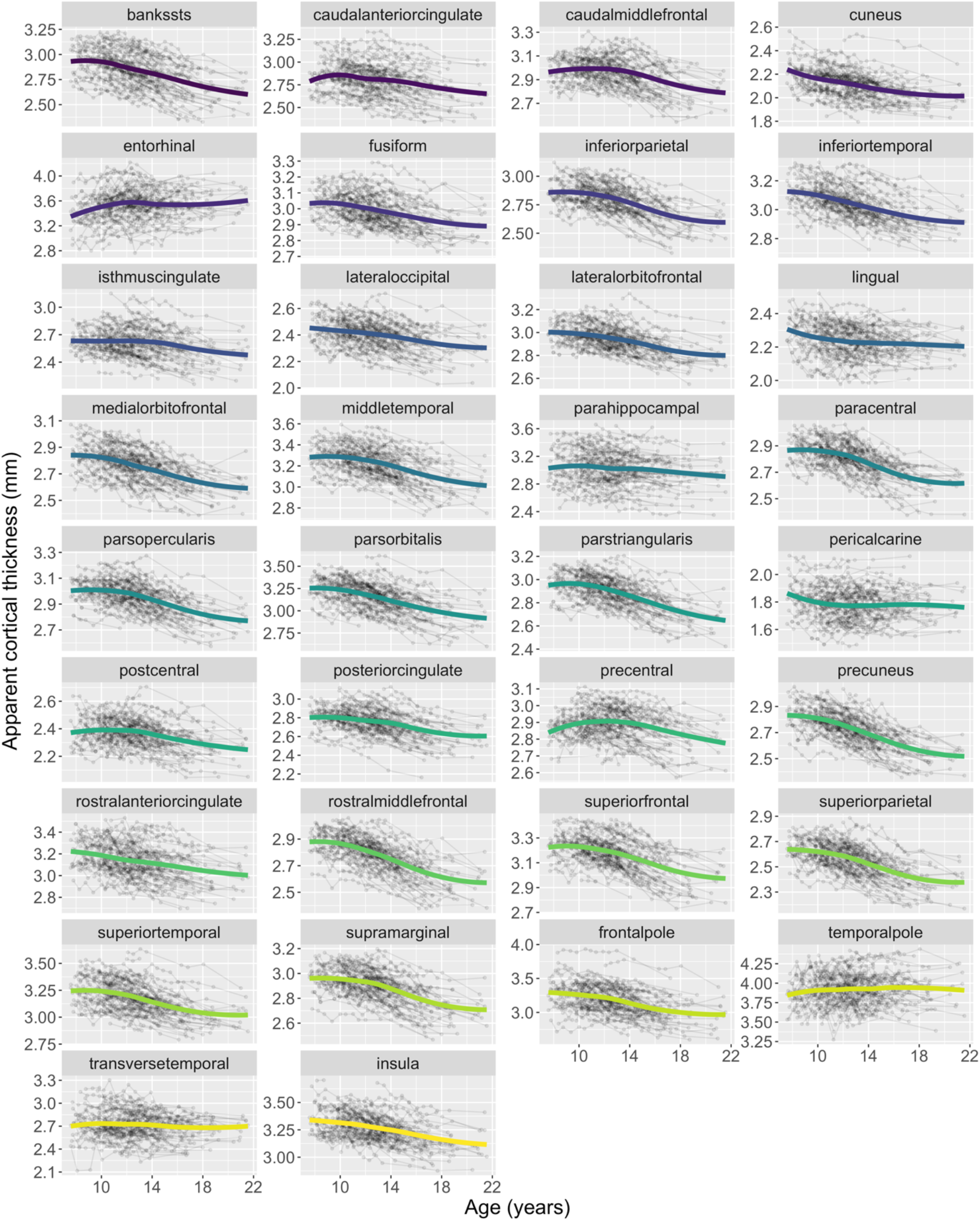
Apparent cortical thickness across brain regions (defined by the Desikan-Killiany atlas) as a function of age. Average and individual trajectories for each participant are shown.

Because the sigmoid is asymptotic, there is no age at which the brain is mature. Instead, the brain develops throughout the age range investigated here (7 – 21 years). Parameters indicated that mean thickness showed high levels in late childhood, with an upper asymptote of 2.95 mm (*SE* = 0.01 mm) and decreasing thereafter to a lower asymptote of 2.62 mm (*SE* = 0.03 mm). Our central parameter of interest, the MCT, was estimated to be 14.36 years (*SE* = 0.28 years). See Table 1. We observed a substantial range of MCTs across individuals, with a minimum and maximum of 12.25 and 19.54 years (Figure 4) and a variance of 1.79 years (*SE* = 0.46 years). Together, 0ur analysis demonstrates that we can estimate a novel, quantitative definition of cortical maturity which is independent of overall thickness and shows substantial differences between people.

**Table 1:** NLMM Parameter estimates for mean cortical thickness

| Parameter | Estimate | SE | CV (%) | *p* |
| --- | --- | --- | --- | --- |
| $A_{Lower}$ | 2.62 | 0.03 | 1.3 | - |
| $b_{Sex(A_{Lower})}$ | -0.07 | 0.04 | 55.1 | 0.035 |
| $b_{Motion(A_{Lower})}$ | 0.00 | <0.01 | 52.0 | 0.027 |
| $A_{Upper}$ | 2.95 | 0.01 | 0.4 | - |
| $b_{Sex(A_{Upper})}$ | -0.06 | 0.02 | 30.5 | 0.001 |
| $MCT$ | 14.36 | 0.28 | 2.0 | - |
| $b_{Sex(MCT)}$ | 1.92 | 0.51 | 26.7 | <0.001 |
| $Hill$ | -6.34 | 0.42 | 6.6 | - |

Note. CV = Coefficient of variation – the ratio of the standard deviation to the mean. Used to assess precision. CV values $\leq 20\%$ are deemed acceptable. SE = standard error, NLMM = Nonlinear Mixed Models. Females were coded as the reference group.

**Figure 4.**
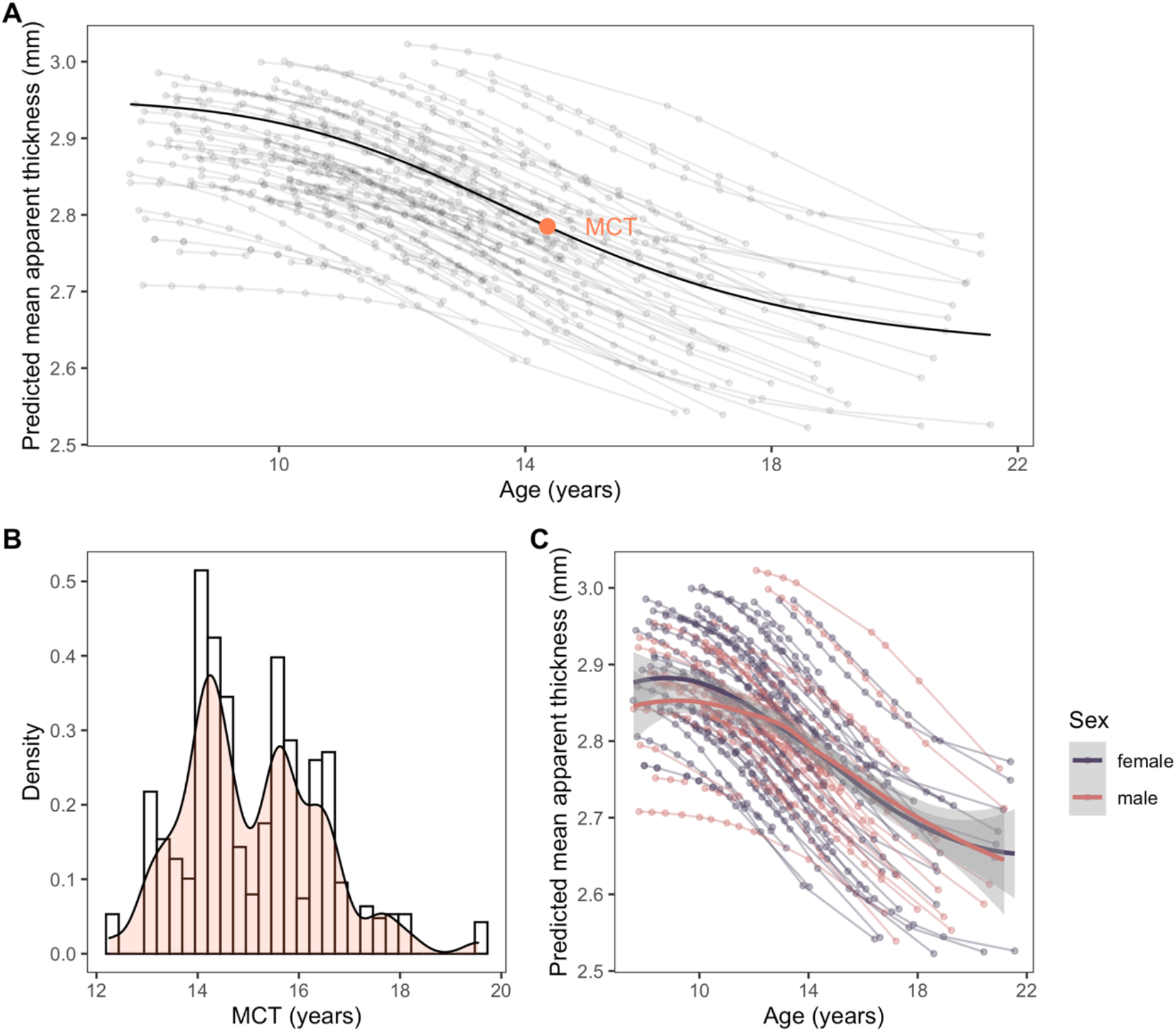
*Panel A*: Predicted apparent mean cortical thickness as a function of age. Estimated individual trajectories and the average trajectory is shown, as well as the average MCT. *Panel B*: Density plot of MCTs showing individual differences in the sample. *Panel C:* Sex differences in the sample. MCT = Midpoint of Cortical Thinning.

### There are substantial individual differences in the MCT

We found that the nonlinear model fit the data well (Supplementary Figure 1). To test whether our model fit better than a simpler alternative, we compared the model fit of the four-parameter logistic nonlinear model to a simple linear model. We found that the four-parameter logistic model fits better than the simple linear model (ΔAIC = -137.22), suggesting that our more complex model is plausible across most regions. Mean cortical thickness was estimated with very good precision for the asymptotes, MCT, and the hill, as indicated by low coefficients of variation (*CV*; Table 1). The MCT was uncorrelated with the upper asymptote, showing that it is independent of the initial thickness of a region (*r* = 0.01, *t*(88) = 0.09, *p* = .925). It was also independent of the mean cortical thickness in early adulthood (*A*_*Lower*_, *r* = -0.13, *t*(88) = -1.22, *p* = .227). This independence means that cross-sectional measurements of cortical thickness will likely not function well as an approximation of cortical maturity. The rate of change (*Hill*) was also not associated with the MCT (*r* = -0.16, *t*(88) = -1.48, *p* = .141). A sensitivity analysis shows that results were very similar when only participants with at least three scans were included (*N* = 84, Supplementary Table 2).

### Significant, but noisy, sex differences in trajectories

There were significant sex differences: Females started with thicker cortices than males (*b* = -0.06, *p* = .001), had thicker corteces in adulthood (*b* = -0.07, *p* = .035) and showed earlier MCT (*b* = 1.92, *p* < .001, Figure 4). However, the *CVs* for these estimates were all well above 20% (Table 1), indicating that these parameters were estimated with low precision. The low precision was likely due to the small number of males in the sample (*N*_*Males*_ = 37).

### MCTs differ across the cortex

To examine the specificity in maturational timing across regions, we estimated the MCT for each of the 34 cortical regions of the Desikan-Killiany atlas independently (Figure 5), again using non-linear mixed models and the four-parameter logistic function. The four parameters were estimated with good precision (*CV* < 20%) for all regions^51^, except for the entorhinal, lingual, pericalcarine, temporal pole, and transverse temporal regions, which were all excluded from subsequent analyses (see Supplementary Tables 3 and 4 for all estimates). For these regions, the logistic may not be a good approximation of the functional relationship between age and cortical thickness. For the pericalcarine, for instance, we find that a linear model fits marginally better than the four-parameter logistic (ΔAIC = 0.50), potentially reflecting the early maturation for this region. The enthorinal is known to be more likely to be affected by eye-movement artifacts and may therefore show a poor model fit.

**Figure 5.**
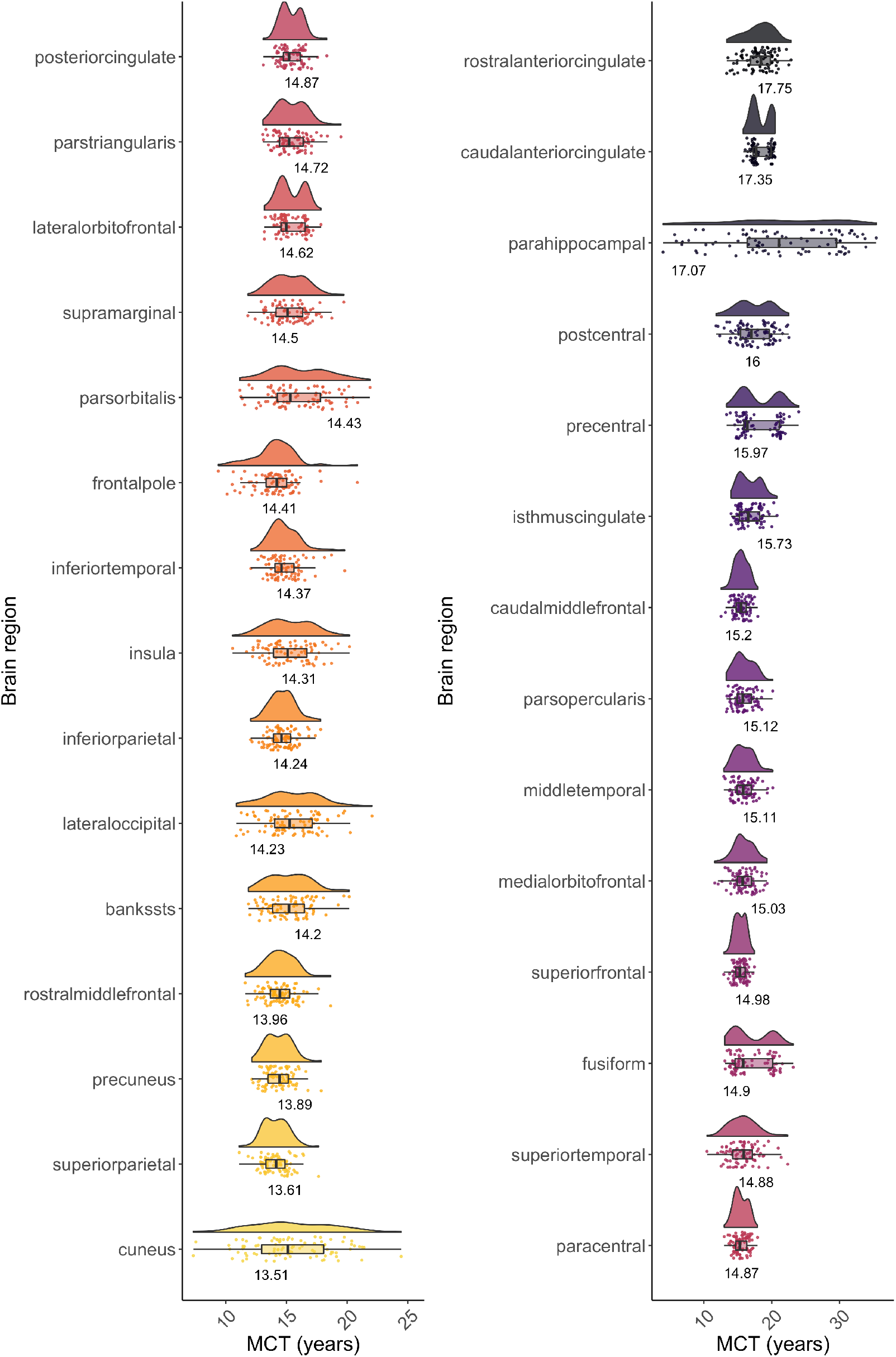
Individual and regional differences in thinning shown in raincloud plots. The estimated kernel density is shown to visualize the distribution of values. The median is shown as the black vertical line within a bar, which itself shows the interquartile range. Black horizontal lines show the 1.5 interquartile range. Values beyond these lines can be considered outliers.

Mean MCTs ranged from 13.51 years for the cuneus to 17.75 years for the rostral anterior cingulate cortex (Figure 5). The brain regions that reached the MCT first were the occipital (cuneus and lateral occipital) and parietal (precuneus, superior and inferior parietal) cortices, the rostral middle frontal cortex, and the cortex of the banks of the superior temporal sulcus (bankssts). The brain regions that reached the MCT last were the caudal and rostral anterior cingulate and parahippocampal cortices, followed by the sensorimotor pre- and postcentral. Moreover, the frontal lobe regions (precentral, caudal middle frontal, rostral middle frontal, superior frontal, and frontal pole) showed the fastest rate of change (i.e., steepest hill), while the slowest rate of change (i.e., flattest hill) was observed in the parahippocampal, cuneus, (caudal and rostral) anterior cingulate, and superior temporal cortices. Most regions showed a relative independence of the MCT, upper and lower asymptote and hill (Figure 6) and low parameter correlations (Supplementary Table 5).

**Figure 6.**
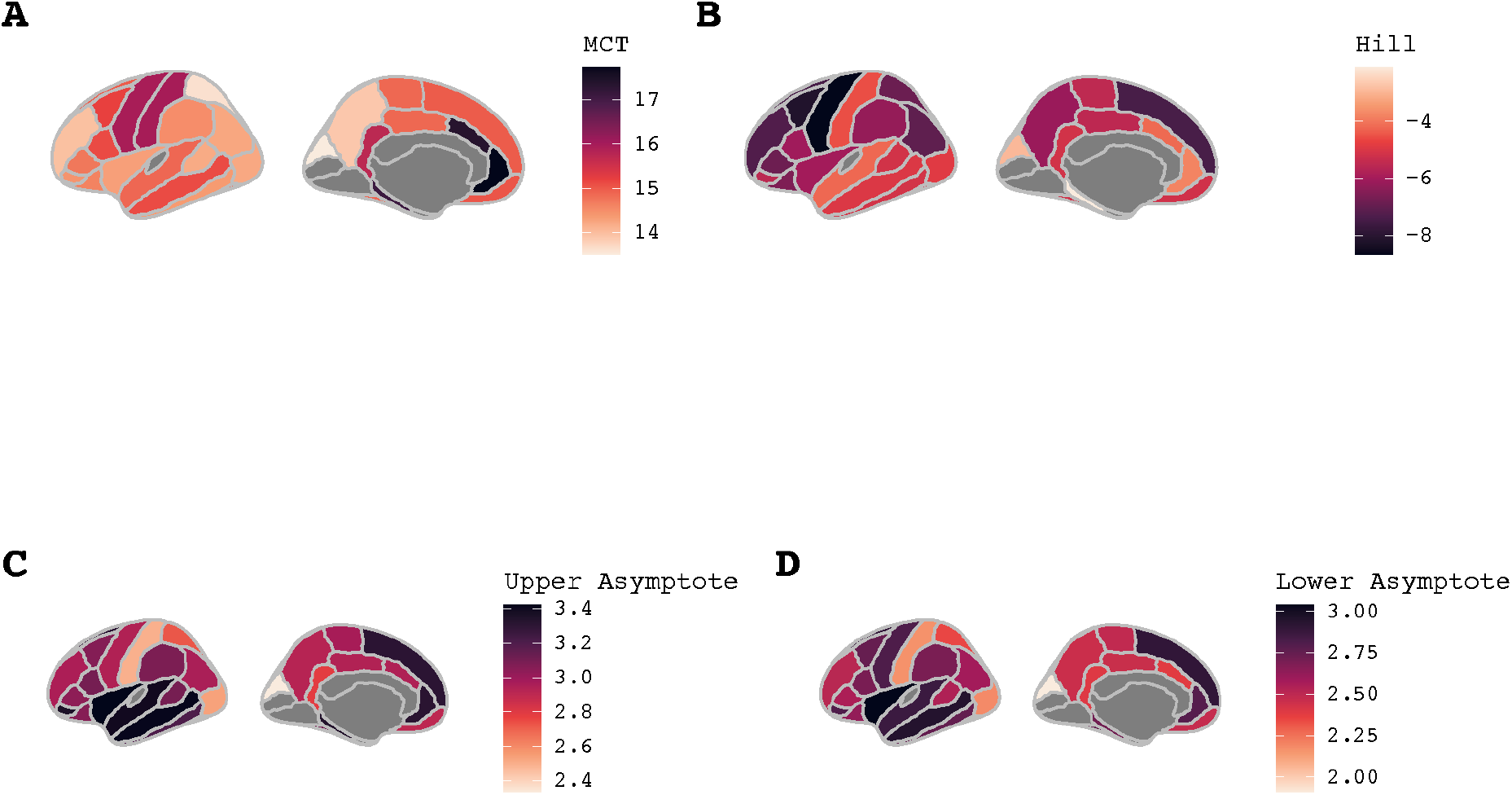
Parameter estimates for the MCT (Panel A), hill (Panel B), upper asymptote (Panel C) and lower asymptote (Panel D) plotted across the cortex. Darker shades reflect higher parameter estimates. Excluded regions are shown in grey.

Next, we implemented a formal quantitative test to examine whether regions differed in their MCTs. To do so, we compared a confirmatory factor model with intercepts for MCTs across regions constrained to equality (reflecting the hypothesis that regions mature at the same approximate age) to a model where they are estimated freely, reflecting the hypothesis that regions mature at distinct ages). Despite the considerable added complexity, we found that allowing MCTs to differ between regions, substantially improved model fit (ΔX^2^(29) = 930.69, *p* < .001) indicating pronounced differences in maturational timing between regions independent of their overall thickness.

In addition to MCT-differences across regions globally, we also examined whether MCTs are linked between some or all regions: In other words, if a person is mature in one region, are they then also more mature in all other regions, or are there clusters of brain regions that covary in their relative maturity? To examine this question, we first examined the fit of a one-factor confirmatory model, testing the hypothesis that a single factor could capture MCTs across the brain. This model fit poorly (*X*^2^(377) = 1236.68, *p* < .001, CFI = 0.787, SRMR = 0.059, RMSEA = 0.159 [0.150, 0.169]). To determine whether a more complex model might fit the data, we used Exploratory Factor Analysis. Eigenvalues of a parallel analysis suggested that a two-factor model yields the best, albeit imperfect, solution (*X*^2^(349) = 633.85, *p* < .001). The two-factor model explained 79.3% of the variance in MCTs. Several central, as well as cingulate, regions loaded strongly onto Factor 1. Parietal regions regions and the rostral middle frontal loaded strongly onto Factor 2 (Figure 7, Supplementary Table 6).

**Figure 7.**
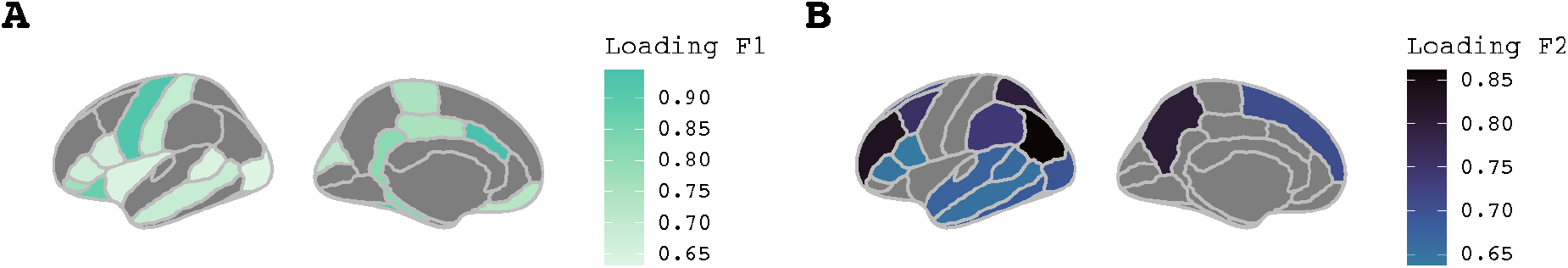
Factor loading for Factor 1 (Panel A) and Factor 2 (Panel B) across the cortex. Only loadings > 0.6 are shown to facilitate interpretation. Darker colours represent higher loadings.

Finally, we explored whether there is a regional ordering in the timing of maturation in line with hypotheses of a posterior-to-anterior gradient across the cortex^7,56^. We correlated inflection ages of each region with the region’s average y-coordinate in MNI space, as contained in mni.y in brainGraph^55^. We found no evidence for a significant correlation between the spatial location and the MCT (*t*(27) = 0.12, *p* = .903, *r* = 0.02, Bayes Factor = 0.35). In an additional exploratory analysis, we also analyzed whether there is a regional ordering in the timing of maturation in line with hypotheses of a dorsal-ventral gradient across the cortex^57^. We correlated inflection ages of each region with the region’s average z-coordinate in MNI space, as contained in mni.z in brainGraph^55^. We found no evidence for a significant correlation between the spatial location and the inflection ages (*t*(19) = -0.65, *p* = .522, *r* = - 0.15, Bayes Factor = 0.45).

Together, these analyses offer new insight into cortical maturation. We demonstrate, for the first time, that it is possible to estimate non-linear maturation independent of overall cortical thickness. Maturational trajectories differed between individuals and cortical regions. The ability to estimate these differences offers a new window into elucidating long-standing debates concerning the speed of maturation, its association with early adversity, and the implications for cognitive and mental health development.

## Discussion

Using longitudinal data of up to 12-waves imaged between late childhood and early adulthood and flexible nonlinear mixed models, we here show that cortical maturity as indexed by the MCT can be separated from other cortical thickness parameters (i.e., asymptotes and slope – hill) and estimated precisely and reliably for each individual and most brain regions. We identified a characteristic, s-shaped, trajectory of cortical thinning: Cortical thickness was high in childhood, followed by decreases in early adolescence, culminating around the age of 14 years, the average MCT. The reduction in cortical thickness then decelerates to level off in late adolescence. This finding is in line with previous studies showing s-shaped reductions in cortical thickness over adolescence^9,12^ and extend the same by providing estimates of upper limits in apparent cortical thickness (2.95 mm), lower limits of apparent cortical thickness (2.62 mm), and, most importantly, an index of maturation: the MCT (average of 14.36 years). This highlights that cortical thinning is protracted and shows rapid changes in adolescence. This period of rapid brain development raises questions about possible sensitive periods in adolescence^58,59^. Future research will be able to show whether periods of structural change confer heightened plasticity in adolescence.

Developmental patterns differed between cortical regions, with the superior parietal and precuneus showing some of the earliest MCTs, around 14 years, and cingulate regions showing some of the latest MCTs, around 17 years. Early maturing regions were found in lateral frontal, and parietal areas, while late-maturing regions were found in temporal and dorsal central areas.

Our finding of an early MCT in several frontal areas is surprising, given previous, mostly cross-sectional studies. There are several possible explanations for this result, which will need to be tested in future research: First, while most frontal regions are usually imaged well, the frontal pole is known to be difficult to image. We therefore advise to interpret the finding for this region with caution. Second, we cannot rule out that data quality issues or model misfit may have affected findings, although quality control procedures, statistical control for in-scanner motion, diagnostic plots and precision estimates do not suggest that this was likely to be the sole explanation. Third, while the scans were evenly distributed across most ages, the interval between waves 11 and 12 was longer, and data was sparser at older ages. This may cause the model fit to be poorer at older ages, although this would be more likely to affect estimation of late maturing, rather than early maturing regions. Fourth, the late maturation suggested by the, mostly, cross-sectional analyses to date, do not replicate in appropriate longitudinal analyses like ours, because of the inherent limitation of cross-sectional data for the examination of developmental patterns. Finally, it may be that cross-sectional analyses have captured aspects of maturation that are independent of the MCT, the time point of fastest thinning. It may be, for instance, that frontal regions asymptote later than other region, even though the peak of thinning happens quite early. Altogether this finding, along with recent work identifying structural brain networks and hubs^60–63^, supports a complex systems account, in which the brain matures in a distributed pattern. Future studies could use complex systems approaches like network models to identify how maturation across regions produces changes in cognition and mental states across development.

We found evidence for pronounced individual differences, with MCTs differing by several years between individuals. This supports the notion that brain maturation is highly variable and invites questions of potential predictors and outcomes. We here investigated sex as a potential predictor of individual differences. There was some evidence for sex differences, with females showing earlier maturation than males, by about 2 years. This supports initial evidence for earlier maturation in girls from older longitudinal studies^46,47^ and may be linked to earlier puberty and different socialization in girls^64,65^. Our estimates of sex differences were relatively noisy, however. Future studies could use nonlinear mixed models in larger cohorts, such as ABCD^28^, to investigate the robustness of sex differences in maturation. Future studies could also investigate other candidate predictors such as environmental influences (e.g., adversity and education) to identify whether these accelerate or decelerate maturation – a yet unsolved conundrum in developmental science^18,19,66^. Investigations of potential outcomes of maturational differences, e.g., cognitive performance or psychiatric diagnosis, would be similarly fascinating, and could eventually include distal effects, such as cognitive and brain changes during ageing^1,67^.

It is worth noting that our analytical approach, using the four-parameter logistic function, depends on the nature of cortical thinning: Other measures of brain maturation (e.g., brain v0lume) likely show different developmental shapes and different maturational timelines, not all of which will be amenable to estimating the MCT. The simple linear decreases, reported for volume changes in some cortical regions, would not allow for a quantitative ‘midpoint.’ However, nonlinear mixed models are extremely versatile and can be fit, in principle, to any functional relationship. This also includes more complex relationships than that captured by the four-parameter logistic fit here, allowing, e.g., for asymmetries in development before and after adolescence. In the future, these nonlinear mixed models can be used to model white matter trajectories and other morphometric measures of cortical and subcortical development to better understand similarities and differences across brain tissues. This will yield a more precise understanding of how changes in grey and white matter work in concert to produce functional changes in the brain and behaviour.

Readers may be interested in understanding the power prerequisites for using nonlinear mixed models. Power in nonlinear mixed models depends on the interaction between several factors, including the number of participants, number of timepoints, spacing between time points, missingness and model complexity^68,69^. While past studies of optimal design in pharmacology indicate that at least three time points may be sufficient to estimate simple nonlinear mixed models^70^, future studies will need to evaluate power for plausible developmental functions and typical designs of neuroimaging studies in detail.

In conclusion, this study shows that apparent cortical thinning in adolescence is s-shaped, with the most rapid changes occurring in mid-adolescence, at around 14 years of age, on average. Further, we show that individuals vary substantially, with up to several years, in the age at which the cortex undergoes most rapid changes. On a practical level, this work shows that high-resolution temporal data, combined with nonlinear modelling approaches, can be used to quantify brain maturation with unprecedented precision. This will allow the field to provide rigorous tests of prominent theoretical models of adolescent development, such as the structural mismatch hypothesis^20,21^ or accelerated maturation hypotheses^17^.

## References

1. Bethlehem, R. A. I. et al. Brain charts for the human lifespan. bioRxiv 2021.06.08.447489 (2021) doi:10.1101/2021.06.08.447489.

2. Fuhrmann, D., Simpson-Kent, I. L., Bathelt, J., The CALM Team & Kievit, R. A. A hierarchical watershed model of fluid intelligence in childhood and adolescence. Cerebral Cortex (2019) doi:10.1093/cercor/bhz091.

3. Humphreys, K. L. et al. Stressful Life Events, ADHD Symptoms, and Brain Structure in Early Adolescence. Journal of Abnormal Child Psychology 47, 421–432 (2019).

4. Mofrad, S. A., Lundervold, A. J., Vik, A. & Lundervold, A. S. Cognitive and MRI trajectories for prediction of Alzheimer’s disease. Scientific Reports 11, 2122 (2021).

5. Bos, M. G. N., Peters, S., van de Kamp, F. C., Crone, E. A. & Tamnes, C. K. Emerging depression in adolescence coincides with accelerated frontal cortical thinning. Journal of Child Psychology and Psychiatry 59, 994–1002 (2018).

6. Shaw, P. et al. Intellectual ability and cortical development in children and adolescents. Nature 440, 676–679 (2006).

7. Squeglia, L. M., Jacobus, J., Sorg, S. F., Jernigan, T. L. & Tapert, S. F. Early adolescent cortical thinning is related to better neuropsychological performance. Journal of the International Neuropsychological Society : JINS 19, 962–70 (2013).

8. Walhovd, K. B., Fjell, A. M., Giedd, J., Dale, A. M. & Brown, T. T. Through Thick and Thin: a Need to Reconcile Contradictory Results on Trajectories in Human Cortical Development. Cerebral Cortex 27, (2017).

9. Tamnes, C. K. et al. Development of the cerebral cortex across adolescence: A multisample study of interrelated longitudinal changes in cortical volume, surface area and thickness. Journal of Neuroscience (2017) doi:10.1523/JNEUROSCI.3302-16.2017.

10. Cardinale, F. et al. Validation of FreeSurfer-Estimated Brain Cortical Thickness: Comparison with Histologic Measurements. Neuroinformatics 12, 535–542 (2014).

11. Furlong, C. et al. Application of stereological methods to estimate post-mortem brain surface area using 3 T MRI. Magnetic resonance imaging 31, 456–465 (2013).

12. Ducharme, S. et al. Trajectories of cortical thickness maturation in normal brain development--The importance of quality control procedures. Neuroimage 125, 267–279 (2016).

13. Jernigan, T. L., Baare, W. F., Stiles, J. & Madsen, K. S. Postnatal brain development: structural imaging of dynamic neurodevelopmental processes. Progress in brain research 189, 77–92 (2011).

14. Natu, V. S. et al. Apparent thinning of human visual cortex during childhood is associated with myelination. Proc Natl Acad Sci USA 116, 20750 (2019).

15. Huttenlocher, P. R. Synaptic density in human frontal cortex - Developmental changes and effects of aging. Brain research 163, 195–205 (1979).

16. Keding, T. J. et al. Differential Patterns of Delayed Emotion Circuit Maturation in Abused Girls With and Without Internalizing Psychopathology. AJP 178, 1026–1036 (2021).

17. Belsky, J. Early-Life Adversity Accelerates Child and Adolescent Development. Curr Dir Psychol Sci 28, 241–246 (2019).

18. Tooley, U. A., Bassett, D. S. & Mackey, A. P. Environmental influences on the pace of brain development. Nature Reviews Neuroscience 22, 372–384 (2021).

19. Colich, N. L., Rosen, M. L., Williams, E. S. & McLaughlin, K. A. Biological aging in childhood and adolescence following experiences of threat and deprivation: A systematic review and meta-analysis. Psychol Bull 146, 721–764 (2020).

20. Steinberg, L. Risk taking in adolescence: what changes, and why? Ann. N. Y. Acad. Sci. 1021, 51–58 (2004).

21. Casey, B. J., Getz, S. & Galván, A. The adolescent brain. Dev Rev 28, 62–77 (2008).

22. Giedd, J. N. et al. Brain development during childhood and adolescence: A longitudinal MRI study. Nature neuroscience 2, 861–863 (1999).

23. Vijayakumar, N. et al. Thinning of the lateral prefrontal cortex during adolescence predicts emotion regulation in females. Soc Cogn Affect Neurosci 9, 1845–1854 (2014).

24. Peper, J. S., Burke, S. M. & Wierenga, L. M. Chapter 3 - Sex differences and brain development during puberty and adolescence. in Handbook of Clinical Neurology (eds. Lanzenberger, R., Kranz, G. S. & Savic, I.) vol. 175 25–54 (Elsevier, 2020).

25. Nunes, A. S. et al. Atypical age-related changes in cortical thickness in autism spectrum disorder. Scientific Reports 10, 11067 (2020).

26. Vijayakumar, N., Mills, K. L., Alexander-Bloch, A., Tamnes, C. K. & Whittle, S. Structural brain development: A review of methodological approaches and best practices. Developmental Cognitive Neuroscience 33, 129–148 (2018).

27. Mills, K. L. et al. Inter-individual variability in structural brain development from late childhood to young adulthood. Neuroimage 242, 118450–118450 (2021).

28. Casey, B. J. et al. The Adolescent Brain Cognitive Development (ABCD) study: Imaging acquisition across 21 sites. Developmental Cognitive Neuroscience 32, 43–54 (2018).

29. Storsve, A. B. et al. Differential longitudinal changes in cortical thickness, surface area and volume across the adult life span: regions of accelerating and decelerating change. J Neurosci 34, 8488–8498 (2014).

30. Schnack, H. G. et al. Changes in Thickness and Surface Area of the Human Cortex and Their Relationship with Intelligence. Cerebral Cortex 25, 1608–1617 (2015).

31. Davidian, M. & Giltinan, D. M. Nonlinear models for repeated measurement data: An overview and update. Journal of Agricultural, Biological, and Environmental Statistics 8, 387 (2003).

32. Madsen, K. S. et al. Maturational trajectories of white matter microstructure underlying the right presupplementary motor area reflect individual improvements in motor response cancellation in children and adolescents. NeuroImage 220, 117105 (2020).

33. Angstmann, S. et al. Microstructural asymmetry of the corticospinal tracts predicts right–left differences in circle drawing skill in right-handed adolescents. Brain Structure and Function 221, 4475–4489 (2016).

34. Gonzalez, M. R. et al. Brain structure associations with phonemic and semantic fluency in typically-developing children. Dev Cogn Neurosci 50, 100982–100982 (2021).

35. Madsen, K. S., Jernigan, T. L., Vestergaard, M., Mortensen, E. L. & Baaré, W. F. C. Neuroticism is linked to microstructural left-right asymmetry of fronto-limbic fibre tracts in adolescents with opposite effects in boys and girls. Neuropsychologia 114, 1–10 (2018).

36. Madsen, K. S. et al. Brain microstructural correlates of visuospatial choice reaction time in children. NeuroImage 58, 1090–1100 (2011).

37. Klarborg, B. et al. Sustained attention is associated with right superior longitudinal fasciculus and superior parietal white matter microstructure in children. Hum Brain Mapp 34, 3216–3232 (2013).

38. Plachti, A. et al. Only females show a stable association between neuroticism and microstructural asymmetry of the cingulum across childhood and adolescence: A longitudinal DTI study. bioRxiv 2021.08.31.458188 (2021) doi:10.1101/2021.08.31.458188.

39. Dale, A. M. & Sereno, M. I. Improved Localizadon of Cortical Activity by Combining EEG and MEG with MRI Cortical Surface Reconstruction: A Linear Approach. Journal of Cognitive Neuroscience 5, 162–176 (1993).

40. Dale, A. M., Fischl, B. & Sereno, M. I. Cortical Surface-Based Analysis: I. Segmentation and Surface Reconstruction. NeuroImage 9, 179–194 (1999).

41. Desikan, R. S. et al. An automated labeling system for subdividing the human cerebral cortex on MRI scans into gyral based regions of interest. NeuroImage 31, 968–980 (2006).

42. Reuter, M., Schmansky, N. J., Rosas, H. D. & Fischl, B. Within-subject template estimation for unbiased longitudinal image analysis. Neuroimage 61, 1402–1418 (2012).

43. Rosen, A. F. G. et al. Quantitative assessment of structural image quality. Neuroimage 169, 407–418 (2018).

44. Comets, E., Lavenu, A. & Lavielle, M. Parameter Estimation in Nonlinear Mixed Effect Models Using saemix, an R Implementation of the SAEM Algorithm. Journal of Statistical Software; Vol 1, Issue 3 (2017) (2017) doi:10.18637/jss.v080.i03.

45. An, H., Landis, J. T., Bailey, A. G., Marron, J. S. & Dittmer, D. P. dr4pl: A Stable Convergence Algorithm for the 4 Parameter Logistic Model. The R Journal 11, 171–190 (2019).

46. Lenroot, R. K. et al. Sexual dimorphism of brain developmental trajectories during childhood and adolescence. NeuroImage 36, 1065–1073 (2007).

47. Wierenga, L. M. et al. A Key Characteristic of Sex Differences in the Developing Brain: Greater Variability in Brain Structure of Boys than Girls. Cerebral Cortex 28, 2741–2751 (2017).

48. Bilger, M. & Manning, W. G. Measuring overfitting and mispecification in nonlinear models. Health, Econometrics and Data Group Working Paper 11, 25 (2011).

49. Bertsimas, D., King, A. & Mazumder, R. BEST SUBSET SELECTION VIA A MODERN OPTIMIZATION LENS. The Annals of Statistics 44, 813–852 (2016).

50. Trevor Hastie, Robert Tibshirani, & Ryan Tibshirani. Best Subset, Forward Stepwise or Lasso? Analysis and Recommendations Based on Extensive Comparisons. Statistical Science 35, 579–592 (2020).

51. Reed, G. F., Lynn, F. & Meade, B. D. Use of coefficient of variation in assessing variability of quantitative assays. Clin Diagn Lab Immunol 9, 1235–1239 (2002).

52. Rosseel, Y. lavaan: An R package for Structural Equation Modeling. Journal of Statistical Software 48, 1–36 (2012).

53. The Personality Project’s Guide to R. http://personality-project.org/r/psych/.

54. Shaw, P. et al. Neurodevelopmental trajectories of the human cerebral cortex. J. Neurosci. 28, 3586–3594 (2008).

55. Watson, C. G. brainGraph: Graph Theory Analysis of Brain MRI Data. (2020).

56. MacPherson, S. E. et al. Processing speed and the relationship between Trail Making Test-B performance, cortical thinning and white matter microstructure in older adults. Cortex 95, 92–103 (2017).

57. Chen Chi-Hua et al. Genetic topography of brain morphology. Proceedings of the National Academy of Sciences 110, 17089–17094 (2013).

58. Fuhrmann, D., Knoll, L. J. & Blakemore, S. J. Adolescence as a sensitive period of brain development. Trends in cognitive sciences (2015) doi:10.1016/j.tics.2015.07.008.

59. Frankenhuis, W. E. & Walasek, N. E. Evolution of sensitive periods: Bridging theory and data. Developmental Cogntitive Neuroscience (in press).

60. Baum, G. L. et al. Modular segregation of structural brain networks supports the development of executive function in youth. Current Biology (2017) doi:10.1016/j.cub.2017.04.051.

61. Baum, G. L. et al. Development of structure–function coupling in human brain networks during youth. Proc Natl Acad Sci USA 117, 771 (2020).

62. Power, J. D., Fair, D. A., Schlaggar, B. L. & Petersen, S. E. The development of human functional brain networks. Neuron 67, 735–748 (2010).

63. Oldham, S. & Fornito, A. The development of brain network hubs. Developmental Cognitive Neuroscience 36, 100607 (2019).

64. Goddings, A.-L., Beltz, A., Peper, J. S., Crone, E. A. & Braams, B. R. Understanding the Role of Puberty in Structural and Functional Development of the Adolescent Brain. Journal of Research on Adolescence 29, 32–53 (2019).

65. Hyde, J. S. Gender Similarities and Differences. Annu. Rev. Psychol. 65, 373–398 (2014).

66. Ferschmann, L., Bos, M., Herting, M., Mills, K. L. & Tamnes, C. K. Contextualizing adolescent structural brain development. (2021) doi:10.31234/osf.io/w8jq6.

67. Tamnes, C. K. et al. Brain development and aging: Overlapping and unique patterns of change. NeuroImage 68, 63–74 (2013).

68. Retout, S., Comets, E., Samson, A. & Mentré, F. Design in nonlinear mixed effects models: optimization using the Fedorov-Wynn algorithm and power of the Wald test for binary covariates. Stat Med 26, 5162–5179 (2007).

69. Roy, A. & Ette, E. I. A pragmatic approach to the design of population pharmacokinetic studies. AAPS J 7, E408–E420 (2005).

70. al-Banna, M. K., Kelman, A. W. & Whiting, B. Experimental design and efficient parameter estimation in population pharmacokinetics. J Pharmacokinet Biopharm 18, 347–360 (1990).

