## Supplementary Material for "The midpoint of cortical thinning between late childhood and early adulthood differs between individuals and brain regions: Evidence from longitudinal modelling in a 12-wave neuroimaging sample"

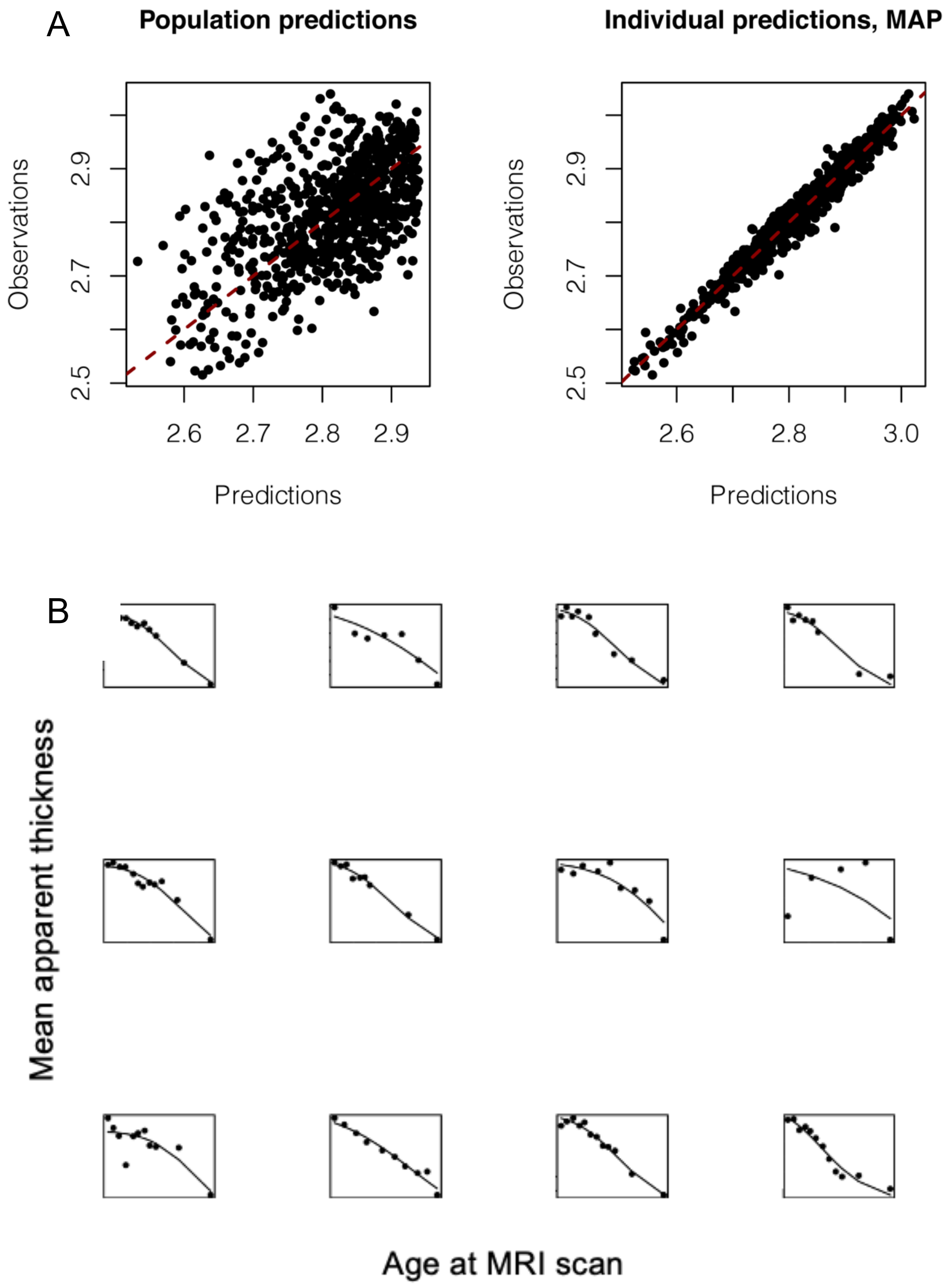


**Supplementary Figure 1**. Diagnostic plots. Panel A: Plots of observed vs. predicted values, both for population prediction and individual predictions. The unity line is shown to aid assessment of deviations. Panel B: Plots showing the individual predictions (solid line) overlaid on the observations (black dots) for a subset of participants in the data set.

**Supplementary Table 1.** Results of the forward model selection procedure used to determine the covariate model.

| **Covariate parameter** | **1st iteration *p*-value** | **2nd iteration *p*-value** | **3rd iteration *p*-value** | **4th iteration *p*-value** | **5th iteration *p*-value** |
|---|---|---|---|---|---|
| sex: lower asymptote | 0.22 | 0.2298 | 0.17484 | 0.03484 | - |
| sex: upper asymptote | 0.0057 | 0.00125 | - | - | - |
| sex: MCT | 0.0035 | - | - | - | - |
| sex: hill | 0.45 | 0.4966 | 0.12118 | 0.06348 | 0.2157 |
| motion: lower asymptote | 0.014 | 0.0598 | 0.01718 | - | - |
| motion: upper asymptote | 0.092 | 0.1026 | 0.08207 | 0.11490 | 0.12 |
| motion: MCT | 0.47 | 0.370 | 0.35137 | 0.21194 | 0.06474 |
| motion: hill | 0.31 | 0.264 | 0.1026 | 0.1755 | 0.40921 |

*Note*. Green cells highlight covariate parameters selected at each iteration.

**Supplementary Table 2**: NLMM Parameter estimates for mean cortical thickness for participants with at least three scans ($N$ = 84)

| **Parameter** | **Estimate** | ***SE*** | ***CV* (%)** | ***p*** |
|---|---|---|---|---|
| $A_{Lower}$ | 2.61 | 0.03 | 1.2 | - |
| $B_{Sex(ALower)}$ | -0.04 | 0.03 | 86.2 | 0.123 |
| $B_{Motion(ALower)}$ | 0.00 | 0.00 | 63.2 | 0.057 |
| $A_{Upper}$ | 2.95 | 0.01 | 0.5 | - |
| $B_{Sex(AUpper)}$ | -0.06 | 0.02 | 31.9 | 0.001 |
| *MCT* | 14.30 | 0.27 | 1.9 | - |
| $B_{Sex(MCT)}$ | 1.49 | 0.46 | 31.3 | 0.001 |
| *Hill* | -6.39 | 0.42 | 6.5 | - |

*Note. CV* = Coefficient of variation – the ratio of the standard deviation to the mean. Used to assess precision. *CV* values ≤ 20% are deemed acceptable. *SE* = standard error, NLMM = Nonlinear Mixed Models. Females were coded as the reference group.

**Supplementary Table 3**. NLMM Parameter estimates for different regions

| Region | $A_{Lower}$ | $b_{Motion}$ *(ALower)* | $b_{Sex}$ *(ALower)* | $A_{Upper}$ | $b_{Sex}$ *(AUpper)* | *MCT* | $b_{Sex}$ *(MCT)* | *Hill* |
|---|---|---|---|---|---|---|---|---|
| bankssts | 2.54 | -0.20 | 0.00 | 3.10 | -0.11 | 14.20 | 2.58 | -5.16 |
| caudalanteriorcingulate | 2.38 | -0.18 | 0.00 | 2.94 | -0.06 | 17.35 | 2.67 | -4.18 |
| caudalmiddlefrontal | 2.75 | -0.05 | 0.00 | 3.06 | -0.04 | 15.20 | 1.08 | -8.15 |
| cuneus | 1.91 | -0.06 | 0.00 | 2.33 | -0.10 | 13.51 | 5.04 | -2.92 |
| entorhinal | 3.55 | 0.08 | 0.00 | 3.27 | -0.13 | 8.61 | -0.90 | -5.13 |
| fusiform | 2.75 | -0.19 | 0.00 | 3.10 | -0.06 | 14.90 | 5.36 | -5.18 |
| inferiorparietal | 2.58 | -0.05 | 0.00 | 2.96 | -0.07 | 14.24 | 1.14 | -7.03 |
| inferiortemporal | 2.75 | 0.00 | 0.00 | 3.20 | -0.04 | 14.37 | 1.25 | -5.13 |
| isthmuscingulate | 2.39 | -0.15 | 0.00 | 2.78 | -0.11 | 15.73 | 2.66 | -5.15 |
| lateraloccipital | 2.18 | -0.11 | 0.00 | 2.53 | -0.03 | 14.23 | 3.31 | -4.86 |
| lateralorbitofrontal | 2.62 | -0.02 | 0.00 | 3.05 | -0.05 | 14.62 | 1.95 | -6.56 |
| lingual | 1.95 | 0.19 | 0.00 | 2.36 | -0.06 | 16.75 | -2.60 | -2.30 |
| medialorbitofrontal | 2.47 | -0.06 | 0.00 | 2.90 | -0.02 | 15.03 | 2.26 | -5.18 |
| middletemporal | 2.94 | -0.14 | 0.00 | 3.41 | -0.05 | 15.11 | 2.13 | -4.97 |
| parahippocampal | 2.71 | -0.16 | 0.00 | 3.33 | -0.26 | 17.07 | 12.91 | -2.09 |
| paracentral | 2.49 | -0.04 | 0.00 | 2.97 | -0.10 | 14.87 | 1.61 | -5.49 |
| parsopercularis | 2.69 | -0.10 | 0.00 | 3.10 | -0.06 | 15.12 | 2.11 | -6.03 |
| parsorbitalis | 2.80 | -0.30 | 0.00 | 3.35 | -0.12 | 14.43 | 4.04 | -5.53 |
| parstriangularis | 2.55 | -0.07 | 0.00 | 3.04 | -0.07 | 14.72 | 1.86 | -6.82 |
| pericalcarine | 1.68 | 0.28 | 0.00 | 1.83 | -0.08 | 14.66 | 14.23 | -2.47 |
| postcentral | 2.16 | -0.11 | 0.00 | 2.49 | -0.08 | 16.00 | 3.81 | -4.56 |
| posteriorcingulate | 2.46 | -0.07 | 0.00 | 2.94 | -0.10 | 14.87 | 1.42 | -5.51 |
| precentral | 2.81 | -0.33 | 0.00 | 2.94 | -0.05 | 15.97 | 5.31 | -8.66 |
| precuneus | 2.46 | -0.01 | 0.00 | 2.92 | -0.09 | 13.89 | 1.26 | -6.15 |
| rostralanteriorcingulate | 2.78 | -0.16 | 0.00 | 3.31 | -0.10 | 17.75 | 1.64 | -3.80 |
| rostralmiddlefrontal | 2.51 | -0.09 | 0.00 | 2.96 | -0.06 | 13.96 | 1.39 | -7.27 |
| superiorfrontal | 2.90 | -0.08 | 0.00 | 3.31 | -0.08 | 14.98 | 1.16 | -7.39 |
| superiorparietal | 2.34 | -0.03 | 0.00 | 2.72 | -0.05 | 13.61 | 1.35 | -6.85 |
| superiortemporal | 2.86 | -0.07 | 0.00 | 3.40 | -0.09 | 14.88 | 2.42 | -4.20 |
| supramarginal | 2.69 | -0.13 | 0.00 | 3.08 | -0.10 | 14.50 | 2.06 | -6.20 |
| frontalpole | 2.73 | 0.05 | 0.00 | 3.37 | -0.11 | 14.41 | -0.53 | -7.18 |
| temporalpole | 3.93 | 0.15 | 0.00 | 3.90 | -0.04 | 26.02 | 5.76 | -1.36 |
| transversetemporal* | 2.52 | -0.14 | 0.00 | 2.90 | -0.12 | 15.78 | 10.86 | -2.56 |
| insula | 3.04 | -0.07 | 0.00 | 3.42 | -0.08 | 14.31 | 2.61 | -5.98 |

*Note:* Regions with *CV*s > 20% in any of the four main parameters of the model are shown in grey and were excluded from further analysis (see Supplementary Table 4). *CV* = Coefficient of Variation, NLMM = Nonlinear Mixed Models. Females were coded as the reference group.
* convergence issues – estimates may not be reliable

**Supplementary Table 4**. NLMM *CV*s (%) for all regions.

| Region | $A_{Lower}$ | $b_{Motion}$ (ALower) | $b_{Sex}$ (ALower) | $A_{Upper}$ | $b_{Sex}$ (AUpper) | *MCT* | $b_{Sex}$ (MCT) | *Hill* |
|---|---|---|---|---|---|---|---|---|
| bankssts | 2.5 | 36.8 | 49.8 | 0.9 | 32.7 | 2.5 | 27.0 | 7.2 |
| caudalanteriorcingulate | 4.1 | 64.9 | 212.3 | 1.0 | 67.1 | 5.2 | 52.6 | 10.6 |
| caudalmiddlefrontal | 1.9 | 99.8 | 55.4 | 0.5 | 62.0 | 1.9 | 42.1 | 7.0 |
| cuneus | 2.5 | 96.3 | 77.0 | 1.6 | 37.6 | 6.3 | 38.2 | 14.6 |
| entorhinal | 1.9 | 69.6 | 207.1 | 2.5 | 135.7 | 11.0 | 154.2 | 26.3 |
| fusiform | 1.6 | 48.7 | 191.7 | 0.5 | 36.0 | 3.0 | 27.3 | 9.5 |
| inferiorparietal | 1.8 | 80.9 | 40.7 | 0.6 | 33.3 | 1.9 | 39.0 | 6.8 |
| inferiortemporal | 1.5 | 1596.0 | 7641.5 | 0.6 | 59.9 | 2.6 | 52.1 | 8.1 |
| isthmuscingulate | 2.7 | 63.2 | 34.2 | 1.0 | 34.5 | 2.8 | 37.9 | 8.5 |
| lateraloccipital | 2.2 | 57.8 | 176.6 | 0.8 | 71.2 | 3.8 | 33.0 | 10.7 |
| lateralorbitofrontal | 1.7 | 217.0 | 42.5 | 0.6 | 54.0 | 2.0 | 31.5 | 8.5 |
| lingual | 3.5 | 29.0 | 81.4 | 1.4 | 58.6 | 11.2 | 117.1 | 21.1 |
| medialorbitofrontal | 2.0 | 107.0 | 207.3 | 0.7 | 150.2 | 3.0 | 40.1 | 9.3 |
| middletemporal | 2.1 | 49.3 | 41.3 | 0.6 | 56.0 | 2.5 | 32.2 | 6.9 |
| parahippocampal | 4.7 | 116.5 | 281.4 | 1.9 | 25.9 | 12.7 | 55.6 | 19.8 |
| paracentral | 1.9 | 145.9 | 217.9 | 0.6 | 24.1 | 2.4 | 42.9 | 8.7 |
| parsopercularis | 1.7 | 57.4 | 61.1 | 0.5 | 37.2 | 2.2 | 29.9 | 7.0 |
| parsorbitalis | 2.3 | 36.8 | 408.0 | 0.7 | 24.0 | 3.0 | 25.6 | 9.0 |
| parstriangularis | 1.7 | 68.3 | 193.7 | 0.5 | 35.2 | 1.8 | 26.3 | 6.7 |
| pericalcarine | 3.5 | 42.0 | 120.8 | 1.2 | 42.8 | 16.0 | 48.8 | 28.4 |
| postcentral | 2.5 | 80.8 | 63.1 | 0.7 | 29.2 | 4.5 | 41.4 | 11.1 |
| posteriorcingulate | 2.2 | 85.1 | 76.1 | 0.7 | 28.4 | 2.0 | 37.9 | 6.7 |
| precentral | 1.9 | 49.6 | 42.0 | 0.4 | 39.1 | 2.6 | 24.7 | 9.7 |
| precuneus | 1.5 | 241.8 | 70.5 | 0.6 | 27.0 | 1.8 | 36.5 | 6.7 |
| rostralanteriorcingulate | 3.4 | 73.9 | 98.3 | 0.8 | 36.6 | 6.8 | 109.5 | 12.6 |
| rostralmiddlefrontal | 1.7 | 46.0 | 105.6 | 0.5 | 40.2 | 1.8 | 30.7 | 6.2 |
| superiorfrontal | 1.9 | 70.2 | 99.7 | 0.5 | 30.3 | 1.7 | 37.8 | 6.6 |
| superiorparietal | 1.7 | 131.5 | 52.6 | 0.6 | 51.9 | 1.9 | 32.6 | 7.3 |
| superiortemporal | 1.6 | 84.9 | 54.9 | 0.8 | 36.9 | 3.4 | 40.8 | 8.6 |
| supramarginal | 1.6 | 34.5 | 39.9 | 0.6 | 23.2 | 2.1 | 26.1 | 6.8 |
| frontalpole | 2.3 | 107.8 | 32.3 | 0.9 | 41.6 | 3.1 | 138.8 | 9.0 |
| temporalpole | 3.4 | 80.6 | 186.8 | 1.0 | 151.8 | 45.2 | 303.8 | 133.6 |
| transversetemporal* | 3.5 | 114.1 | 70.6 | 1.9 | 48.3 | 11.7 | 48.1 | 23.6 |
| insula | 1.6 | 79.7 | 108.9 | 0.7 | 38.6 | 2.9 | 34.0 | 10.0 |

*Note.* Regions with *CV*s > 20% in any of the four main parameters of the model ($A_{Upper}$, $A_{Lower}$, *MCT*, *Hill*) are shown in grey and were excluded from further analysis. *CV* = Coefficient of Variation, NLMM = Nonlinear Mixed Models. Females were coded as the reference group. * convergence issues – estimates may not be reliable

**Supplementary Table 5**: Correlations of NLMM Parameters with the MCT across regions

| **Region** | ***r* (MCT & upper asymptote)** | ***r* (MCT & lower asymptote)** | ***r* (MCT& hill)** |
|---|---|---|---|
| bankssts | 0.01 | -0.28 | -0.18 |
| caudalanteriorcingulate | -0.06 | -0.31 | -0.05 |
| caudalmiddlefrontal | 0.24 | 0.19 | -0.23 |
| cuneus | -0.07 | -0.23 | -0.09 |
| fusiform | -0.19 | -0.69 | 0.08 |
| inferiorparietal | 0.11 | 0.15 | -0.18 |
| inferiortemporal | 0.17 | 0.39 | -0.28 |
| isthmuscingulate | -0.10 | -0.35 | -0.06 |
| lateraloccipital | 0.11 | -0.20 | -0.16 |
| lateralorbitofrontal | -0.12 | 0.09 | -0.01 |
| medialorbitofrontal | 0.17 | -0.01 | -0.08 |
| middletemporal | 0.07 | -0.20 | -0.19 |
| parahippocampal | -0.23 | -0.07 | -0.17 |
| paracentral | -0.13 | 0.13 | -0.12 |
| parsopercularis | 0.01 | -0.24 | -0.22 |
| parsorbitalis | -0.21 | -0.59 | -0.05 |
| parstriangularis | 0.01 | -0.06 | -0.27 |
| postcentral | -0.05 | -0.41 | -0.19 |
| posteriorcingulate | -0.11 | 0.04 | -0.06 |
| precentral | -0.17 | -0.82 | 0.16 |
| precuneus | 0.05 | 0.23 | -0.17 |
| rostralanteriorcingulate | 0.21 | 0.11 | -0.19 |
| rostralmiddlefrontal | 0.05 | 0.02 | 0.00 |
| superiorfrontal | 0.02 | 0.04 | -0.24 |
| superiorparietal | 0.22 | 0.19 | -0.21 |
| superiortemporal | 0.20 | -0.21 | -0.22 |
| supramarginal | -0.04 | -0.30 | -0.16 |
| frontalpole | 0.39 | 0.08 | -0.07 |
| insula | -0.08 | -0.02 | -0.07 |
| bankssts | 0.00 | -0.12 | -0.17 |

**Supplementary Table 6**. Factor loadings for an Exploratory Factor Analysis of the MCT across regions.

| **Region** | **Loadings Factor 1** | **Loadings Factor2** |
|---|---|---|
| bankssts | 0.64 | 0.66 |
| caudalanteriorcingulate | 0.95 | 0.24 |
| caudalmiddlefrontal | 0.46 | 0.76 |
| cuneus | 0.71 | 0.46 |
| fusiform | 0.91 | 0.36 |
| inferiorparietal | 0.44 | 0.86 |
| inferiortemporal | 0.47 | 0.69 |
| isthmuscingulate | 0.81 | 0.45 |
| lateraloccipital | 0.63 | 0.69 |
| lateralorbitofrontal | 0.87 | 0.30 |
| medialorbitofrontal | 0.73 | 0.40 |
| middletemporal | 0.68 | 0.65 |
| parahippocampal | 0.82 | 0.19 |
| paracentral | 0.74 | 0.55 |
| parsopercularis | 0.66 | 0.64 |
| parsorbitalis | 0.77 | 0.50 |
| parstriangularis | 0.67 | 0.65 |
| postcentral | 0.69 | 0.55 |
| posteriorcingulate | 0.75 | 0.51 |
| precentral | 0.92 | 0.31 |
| precuneus | 0.48 | 0.80 |
| rostralanteriorcingulate | 0.38 | 0.37 |
| rostralmiddlefrontal | 0.45 | 0.83 |
| superiorfrontal | 0.59 | 0.71 |
| superiorparietal | 0.49 | 0.80 |
| superiortemporal | 0.52 | 0.67 |
| supramarginal | 0.59 | 0.74 |
| frontalpole | -0.19 | 0.41 |
| insula | 0.64 | 0.55 |
